# Gene III^®^ L-Ergothioneine Ameliorates Exercise-Induced Fatigue by Attenuating Oxidative Stress, Inflammation, and Modulating the AMPK/PGC-1α Signaling Pathway

**DOI:** 10.64898/2026.02.19.706732

**Authors:** Wei Ding, Juan Cao, Cong Guo, Wei Liu, Xu Li, Guohua Xiao

**Author notes:** Correspondence: Guohua Xiao.

## Abstract

**Background:** Exercise-induced fatigue is a complex physiological phenomenon involving oxidative stress, inflammation, and metabolic disturbances. Ergothioneine (EGT), a naturally occurring amino acid with potent antioxidant properties, has garnered interest for its potential health benefits. This study aimed to evaluate the anti-fatigue effects of Gene III^**®**^ EGT in a mouse model of exhaustive exercise and to elucidate its underlying mechanisms.

**Methods:** Male C57BL/6 mice were randomly divided into five groups: a control group (CTL), low-dose EGT (EGT-L, 10 mg/kg), medium-dose EGT (EGT-M, 30 mg/kg), high-dose EGT (EGT-H, 50 mg/kg), and a positive control group (Coenzyme Q10, 50 mg/kg). Mice were subjected to a 4-week treadmill training protocol, followed by an exhaustive running test. We measured exercise performance and collected blood and skeletal muscle samples at multiple time points to assess biochemical markers, inflammatory cytokines, antioxidant status, and key signaling proteins.

**Results:** Gene III^**®**^ EGT supplementation, particularly at medium and high doses, significantly extended the time to exhaustion and running distance. Compared to the control group, EGT treatment significantly reduced post-exercise levels of lactic acid (LA), lactate dehydrogenase (LDH), and blood urea nitrogen (BUN). Furthermore, Gene III^**®**^ EGT suppressed the exercise-induced increase in pro-inflammatory cytokines, including IL-1β, IL-6, and TNF-α. The anti-fatigue effect of EGT was also associated with a reduction in malondialdehyde (MDA) and an increase in the activities of superoxide dismutase (SOD) and glutathione peroxidase (GSH-Px). Mechanistically, EGT promoted the phosphorylation of AMP-activated protein kinase (AMPK) and the expression of peroxisome proliferator-activated receptor-gamma coactivator-1 alpha (PGC-1α) in skeletal muscle, while also increasing the Bcl-2/Bax ratio, suggesting enhanced mitochondrial biogenesis and reduced apoptosis.

**Conclusions:** Our findings demonstrate that Gene III^**®**^ EGT effectively enhances exercise performance and alleviates fatigue. The underlying mechanisms involve the mitigation of oxidative stress and inflammation, as well as the activation of the AMPK/PGC-1α signaling pathway to promote mitochondrial function and cellular protection. These results highlight the potential of Gene III^**®**^ EGT as a nutritional supplement for combating exercise-induced fatigue.

## 1. Introduction

Exercise-induced fatigue is a complex physiological state characterized by a decline in muscle performance and the inability to maintain a desired exercise intensity [1]. It is a common experience for athletes and individuals engaged in strenuous physical activity, and can significantly impact training quality and competitive performance [2]. The pathogenesis of fatigue is multifactorial, involving the depletion of energy reserves, accumulation of metabolic byproducts such as lactic acid, and disturbances in the central and peripheral nervous systems [3]. Among these factors, oxidative stress and inflammation have been identified as key contributors to the development of fatigue [4].

Intense physical exercise leads to a substantial increase in oxygen consumption, which can overwhelm the body’s antioxidant defense systems and result in the overproduction of reactive oxygen species (ROS) [5]. This state of oxidative stress can cause damage to cellular components, including lipids, proteins, and DNA, leading to muscle damage, impaired muscle function, and inflammation [6]. The inflammatory response, characterized by the release of pro-inflammatory cytokines such as interleukin-1β (IL-1β), interleukin-6 (IL-6), and tumor necrosis factor-alpha (TNF-α), further exacerbates muscle damage and contributes to the sensation of fatigue [7]. Therefore, strategies aimed at mitigating oxidative stress and inflammation are considered promising for the prevention and management of exercise-induced fatigue.

Ergothioneine (EGT) is a naturally occurring, sulfur-containing amino acid derivative of histidine, found in various dietary sources, particularly in mushrooms [8]. Humans and other mammals cannot synthesize EGT and must obtain it from their diet [9]. EGT is actively taken up and accumulated in tissues and cells, especially those subjected to high levels of oxidative stress, such as erythrocytes, bone marrow, and the central nervous system, via the specific transporter OCTN1 (organic cation/carnitine transporter novel type 1) [10]. EGT is a potent antioxidant with a unique ability to scavenge a wide range of ROS and reactive nitrogen species (RNS) [11], chelate metal ions [12], and protect cellular components from oxidative damage [13]. Recent studies have suggested that EGT may have various health benefits, including neuroprotective, anti-inflammatory, and anti-aging effects [14]. Given its powerful antioxidant properties, EGT is a promising candidate for combating exercise-induced fatigue.

The mechanisms of EGT’s antioxidant action are multifaceted. EGT can not only prevent the formation of free radicals such as hydroxyl radicals (OH•), but also directly scavenge free radicals and reactive oxygen species (ROS) such as hypochlorite acid (HClO) and peroxynitrite (ONOO‒) [11]. Additionally, EGT can interact with other natural antioxidant defense systems in the body, such as activating intracellular antioxidant pathways involving MAPKs and regulating the levels of peroxidases and antioxidant enzymes like superoxide dismutases [9]. Furthermore, EGT chelates a variety of divalent metal cations (e.g., Fe, Cu, Zn, Ni, and Co), forming redox-inactive ergothioneine-metal complexes which constrain the reactivity of metal ions [12].

Coenzyme Q10 (CoQ10) is a well-known antioxidant and a crucial component of the mitochondrial electron transport chain, playing a vital role in cellular energy production. It has been widely used as a nutritional supplement to improve exercise performance and reduce fatigue [15]. Therefore, in this study, we included CoQ10 as a positive control to benchmark the anti-fatigue effects of EGT.

Although the antioxidant properties of EGT are well-established, its specific effects on exercise-induced fatigue and the underlying mechanisms remain largely unexplored. A recent study reported that ergothioneine supplementation improved aerobic performance in mice without impairing early muscle recovery signaling [16], but the detailed mechanisms involving mitochondrial biogenesis and energy metabolism pathways have not been fully elucidated.

This study was designed to investigate the efficacy of EGT supplementation in a mouse model of exhaustive exercise. We hypothesized that Gene III^**®**^ EGT would improve exercise performance by attenuating oxidative stress and inflammation. Furthermore, we aimed to explore the molecular mechanisms by which Gene III^**®**^ EGT exerts its anti-fatigue effects, with a particular focus on the AMPK/ PGC-1α signaling pathway, a key regulator of mitochondrial biogenesis and energy metabolism [17-18].

## 2. Materials and Methods

### 2.1 Animals

Male C57BL/6 mice, 8 weeks old, were purchased from Jiangsu Jicui Yaokang Biotechnology Co., Ltd. (Animal Production License No: SCXK(Su)2020-0009). All mice were housed in a specific pathogen-free (SPF) environment at a controlled temperature (24–28 °C), relative humidity (50–70%), and a 12-hour light/dark cycle. The animals had free access to standard chow and water. All experimental procedures were conducted in strict accordance with the guidelines of the Animal Ethics Committee (Ethics Approval No: YSL-202511088). After a quarantine and acclimatization period, during which their general health, food and water intake, and body weight were monitored, only healthy mice were selected for the subsequent experiments.

### 2.2 Experimental Design

A total of 50 mice were randomly divided into five groups (n = 10 per group): (1) Control Group (CTL) received an equivalent volume of saline and corn oil by gavage plus treadmill modeling; (2) Low-Dose EGT Group (EGT-L) received 10 mg/kg of EGT daily by gavage plus treadmill modeling; (3) Medium-Dose EGT Group (EGT-M) received 30 mg/kg of EGT daily by gavage plus treadmill modeling; (4) High-Dose EGT Group (EGT-H) received 50 mg/kg of EGT daily by gavage plus treadmill modeling; and (5) Coenzyme Q10 Positive Control Group (Q10) received 50 mg/kg of CoQ10 daily by gavage plus treadmill modeling. All other conditions, including housing, treadmill training, and testing procedures, were kept identical for all groups to ensure the comparability of the results.

### 2.3 Preparation of EGT and CoQ10

Gene III^**®**^ EGT (purity ≥99.99%) was provided by Jiangsu Gene III Biological Technology Co., Ltd. It was dissolved in physiological saline to prepare stock solutions for the low, medium, and high dose groups. Coenzyme Q10 (purity ≥98%, Aladdin Reagent) was dissolved in corn oil with the aid of ultrasonication. To control for solvent effects, a single-variable principle was applied: the CTL and EGT groups received a supplementary gavage of corn oil (100 μL), while the Q10 group received a supplementary gavage of saline (100 μL). All solutions were prepared fresh and administered daily by oral gavage based on the body weight of each mouse.

### 2.4 Exercise Fatigue Model

#### 2.4.1. Endurance Training Phase

Mice were subjected to a 4-week adaptive endurance training program on a treadmill (SA101C, Jiangsu Saiangsi Biotechnology Co., Ltd.). The training was conducted 5 days a week, with a gradual increase in intensity: Week 1 consisted of 10 m/min at 0° incline for 30 min/day; Week 2 consisted of 12 m/min at 5° incline for 45 min/day; Week 3 consisted of 14 m/min at 5° incline for 60 min/day; and Week 4 consisted of 16 m/min at 5° incline for 60 min/day. During this period, mice received their respective daily treatments.

#### 2.4.2 Exhaustive Endurance Test

After the 4-week training period, all mice underwent an exhaustive endurance test. The test started with a 10-minute warm-up at 14 m/min, after which the speed was increased to 16 m/min with a 5° incline. The mice ran until exhaustion, and the time to exhaustion was recorded. Exhaustion was determined when a mouse was unable to return to the treadmill belt within 10 seconds after falling back onto the shock grid three consecutive times.

### 2.5 Sample Collection and Biochemical Analysis

Blood and skeletal muscle tissue samples were collected at baseline (before the exhaustive run) and at 0, 3, 24, and 48 hours post-exhaustion. Blood samples were allowed to clot at 4 °C and then centrifuged to obtain serum, which was stored at −80 °C for subsequent analysis. Skeletal muscle (gastrocnemius) was excised, and portions were either fixed for histology or snap-frozen in liquid nitrogen and stored at −80 °C for molecular analysis. Serum levels of lactic acid (LA), lactate dehydrogenase (LDH), and blood urea nitrogen (BUN) were measured using commercial assay kits (Jiangsu Edison Biotechnology Co., Ltd., China) according to the manufacturer’s instructions.

### 2.6 Inflammatory Cytokine and Antioxidant Enzyme Assays

Serum concentrations of IL-1β, IL-6, and TNF-α were determined using ELISA kits (Hunan Aifang Biotechnology Co., Ltd., China). The activities of superoxide dismutase (SOD), glutathione peroxidase (GSH-Px), and the concentration of malondialdehyde (MDA) in the serum were measured using corresponding commercial kits (Jiangsu Edison Biotechnology Co., Ltd., China). All procedures were performed following the manufacturers’ protocols.

### 2.7 Histological Analysis

Gastrocnemius muscle samples were fixed in 4% paraformaldehyde, embedded in paraffin, and sectioned. The sections were stained with Periodic Acid-Schiff (PAS) to visualize glycogen content and assess muscle tissue morphology.

### 2.8 Real-Time Quantitative PCR (RT-qPCR)

Total RNA was extracted from skeletal muscle tissue using Trizol reagent. cDNA was synthesized using a reverse transcription kit. RT-qPCR was performed to measure the mRNA expression levels of Bax and Bcl-2, with 36B4 used as the internal reference gene. The relative gene expression was calculated using the 2^−ΔΔCt method.

### 2.9 Western Blot Analysis

Total protein was extracted from skeletal muscle tissue. Protein concentrations were determined using a BCA protein assay kit. Equal amounts of protein were separated by SDS-PAGE and transferred to PVDF membranes. The membranes were incubated with primary antibodies against p-AMPK, AMPK, PGC-1α, and GAPDH, followed by incubation with HRP-conjugated secondary antibodies. Protein bands were visualized using an ECL detection system, and GAPDH was used as a loading control.

### 2.10 Statistical Analysis

All data are presented as the mean ± standard deviation (SD). Statistical analysis was performed using one-way analysis of variance (ANOVA) followed by Tukey’s post hoc test for multiple comparisons. A p-value of less than 0.05 was considered statistically significant. All statistical analyses were performed using GraphPad Prism software.

## 3. Results

### 3.1 Gene III^®^ EGT Supplementation Improves Exercise Performance in Mice

EGT supplementation significantly improved the exercise performance of the mice. Compared to the CTL group, the running distance and time to exhaustion were significantly increased in all EGT-treated groups (p < 0.01). The EGT-M and EGT-H groups showed the most significant improvements, with increases in running distance of 103.04% and 98.18%, respectively, and increases in time to exhaustion of 100.07% and 99.01%, respectively. The performance of the EGT-treated groups was comparable to that of the Q10 positive control group. Gene III^**®**^ EGT improved exhaustive exercise endurance across the 10–50 mg/kg dose range, with the medium- and high-dose groups exhibiting significantly greater improvements (P < 0.01)

### 3.2 Gene III^®^ EGT Attenuates Exercise-Induced Metabolic Stress

As shown in the serum lactic acid results (Figure 2A–E), there were no statistically significant differences between groups before training (A). Immediately after training (0 h) (B), lactic acid levels in the Gene III^**®**^ EGT dose groups (EGT-L, EGT-M, EGT-H) and the Q10 group were significantly lower than in the CTL group, decreasing by 10.61%, 10.93%, 15.42%, and 16.23% compared to the CTL group, respectively, suggesting that both GENEIII ergothioneine and the positive control can reduce early lactic acid accumulation after exhaustive exercise. At 3 h after training (C), the EGT-L, EGT-M, and EGT-H groups remained significantly lower than the CTL group, and the Q10 group showed an even more significant reduction, with decreases of 3.63%, 1.86%, 1.16%, and 11.11%, respectively, indicating that lactate clearance and metabolic load relief were more adequate in the intervention groups during the early post-exercise recovery phase. By 24 h after training (D), only the positive control Q10 group was lower than the CTL group by 3.49%, and differences between groups were not significant at 48 h (E), indicating that lactic acid, as an acute metabolic indicator, mainly reflects differences in the early post-exercise window.

**Figure 1.**
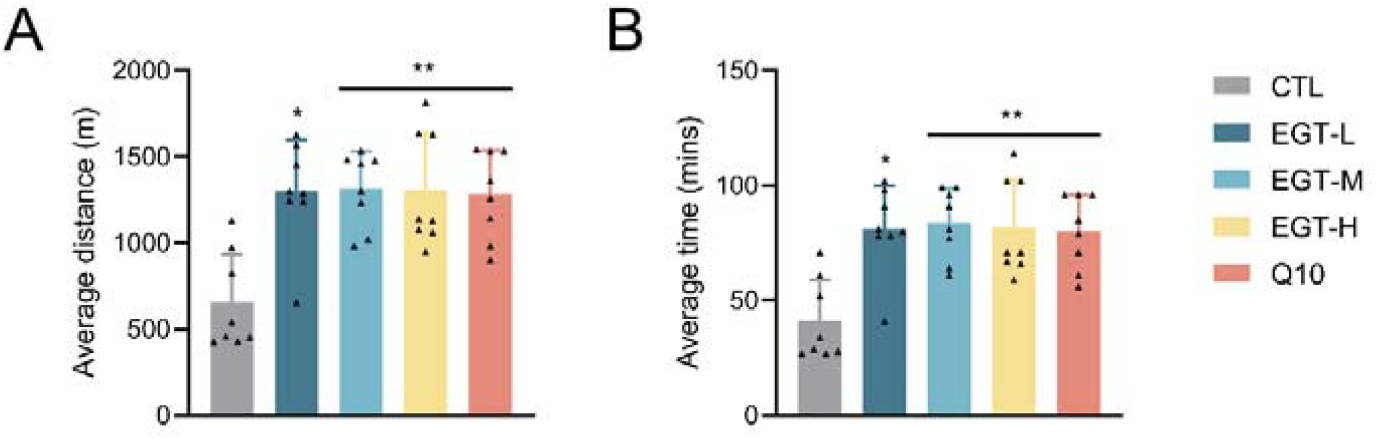
Effects of Gene III^**®**^ EGT on exhaustive exercise performance in mice. (A) Mean running distance (m); (B) Mean running time (min). CTL: vehicle control group; EGT-L, EGT-M, and EGT-H: Gene III**®** EGT groups at 10, 30, and 50 mg/kg, respectively; Q10: Coenzyme Q10 (50 mg/kg) positive control group. Data are expressed as mean $\pm$ SD (n = 10).

**Figure 2.**
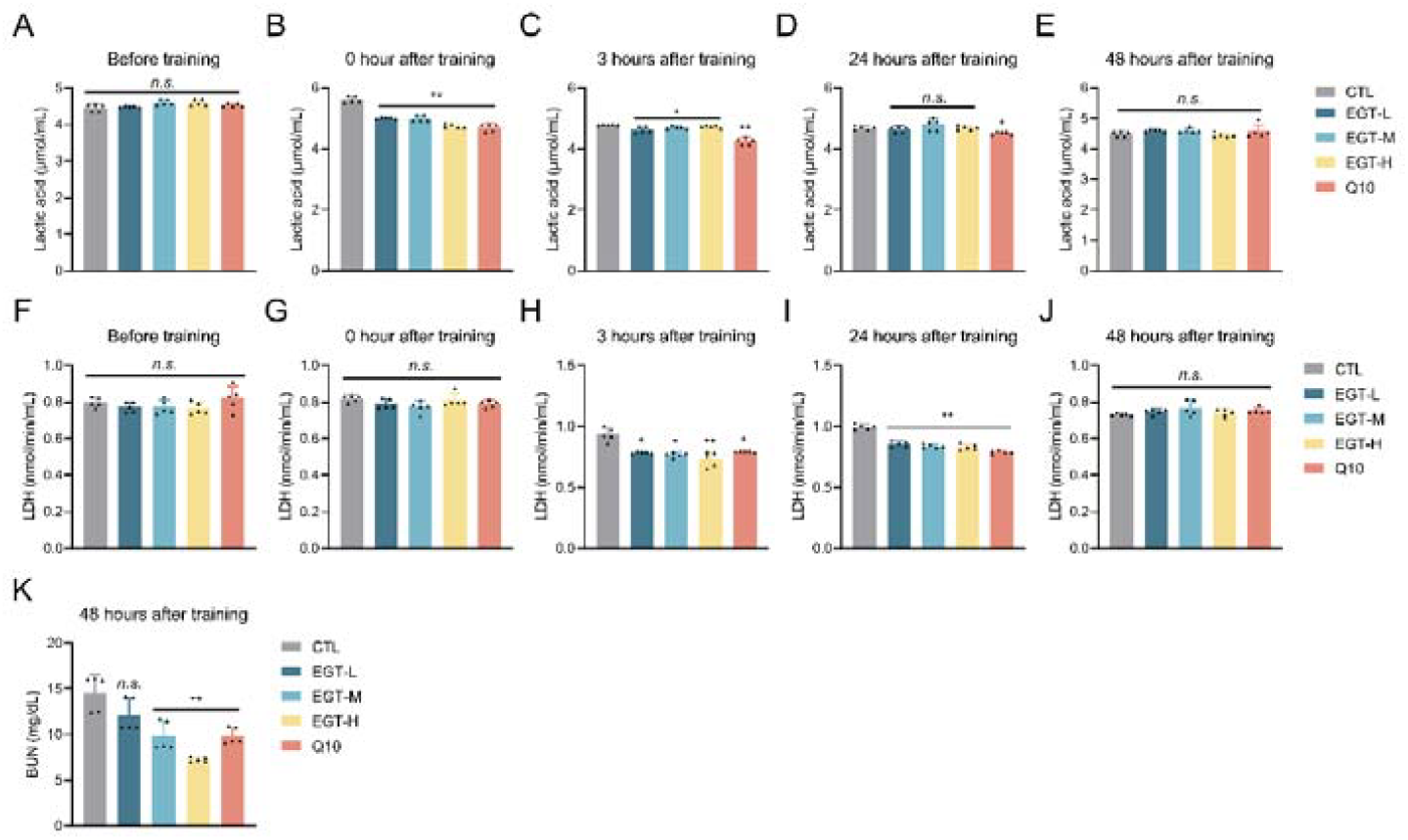
Effects of Gene III^**®**^ EGT on the dynamic changes in serum markers in mice following the exhaustive exercise test. (A–E) Serum lactic acid (μmol/mL): pre-training and 0 h, 3 h, 24h, and 48 h post-training; (F–J) Serum lactate dehydrogenase (LDH, nmol/min/mL): pre-training and 0 h, 3 h, 24 h, and 48 h post-training; (K) Blood urea nitrogen (BUN, mg/dL): 48 h post-training. The initial sample size was 10 animals per group; for analysis, serum samples within the same group were pooled in pairs, resulting in 5 pooled samples per group for detection and statistical analysis (data points in the figure represent pooled samples). CTL: control group; EGT-L, EGT-M, and EGT-H: Gene III^**®**^ EGT groups at 10, 30, and 50 mg/kg, respectively; Q10: Coenzyme Q10 (50 mg/kg) positive control group. Data are expressed as mean ± SD.

As shown in the serum lactate dehydrogenase (LDH) results (Figure 2F–J), differences between groups were not significant before training (F) and at 0 h after training (G). At 3 h after training (H), LDH levels in the EGT-L, EGT-M, and Q10 groups were significantly lower than in the CTL group, with the EGT-H group showing a more significant reduction, decreasing by 16.23%, 17.89%, 21.82%, and 15.31%, respectively. At 24 h after training (I), LDH levels in the EGT-L, EGT-M, EGT-H, and Q10 groups were significantly lower than in the CTL group by 14.14%, 15.89%, 16.76%, and 21.22%, respectively, suggesting that Gene III^**®**^ EGT can effectively reduce serum LDH levels during the 3–24 h post-exercise phase, which may be related to alleviating exercise-induced muscle cell injury, reducing alterations in cell membrane permeability, or promoting tissue repair. By 48 h (J), differences between groups were not significant, consistent with the pattern of acute injury-related serum enzymatic indicators gradually returning to baseline during recovery.

As shown in the blood urea nitrogen (BUN) results (Figure 2K), detection at 48 h after training showed that the EGT-M, EGT-H, and Q10 groups were significantly lower than the CTL group, decreasing by 32.54%, 50.15%, and 32.38% compared to the CTL group, respectively, while the difference in the EGT-L group was not significant. This suggests that medium and high doses of Gene III^**®**^ EGT can reduce the nitrogenous metabolic burden during the recovery period, potentially reflecting a decrease in protein catabolic pressure or an improvement in the overall recovery state after exhaustive exercise.

In summary, Gene III^**®**^ EGT intervention following exhaustive exercise demonstrates a comprehensive ameliorative effect on post-exercise metabolic stress, muscle injury-related changes, and recovery load by reducing early lactic acid levels, lowering LDH levels during the 3–24 h phase, and decreasing BUN during the recovery period. Furthermore, the medium and high doses showed more distinct efficacy on certain recovery indicators.

### 2.2 Gene III^®^ EGT Suppresses Exercise-Induced Inflammation

As shown in the serum IL-1β results (Figure 3A–E), before training (A), the Q10 group exhibited significantly lower levels compared to the CTL group, while the EGT-M and EGT-H groups showed even more significant reductions relative to the CTL group. Specifically, these three groups were significantly lower than the CTL group by 11.26%, 16.66%, and 15.65%, respectively, whereas the EGT-L group showed no significant difference. Immediately after training (0 h; B) and at 3 h post-training (C), both the EGT-H and Q10 groups were significantly lower than the CTL group. At 0 h, they decreased by 4.45% and 0.97%, respectively, compared to the CTL group; at 3 h, they decreased by 8.05% and 12.44%, respectively. This suggests that high-dose Gene III^**®**^ EGT and the positive control Q10 can reduce IL-1β levels during the early phase of the post-exercise inflammatory response. At 24 h post-training (D), the Q10 group remained 10.68% lower than the CTL group, while differences in other groups were not significant. At 48 h (E), there were no significant differences among the groups.

**Figure 3.**
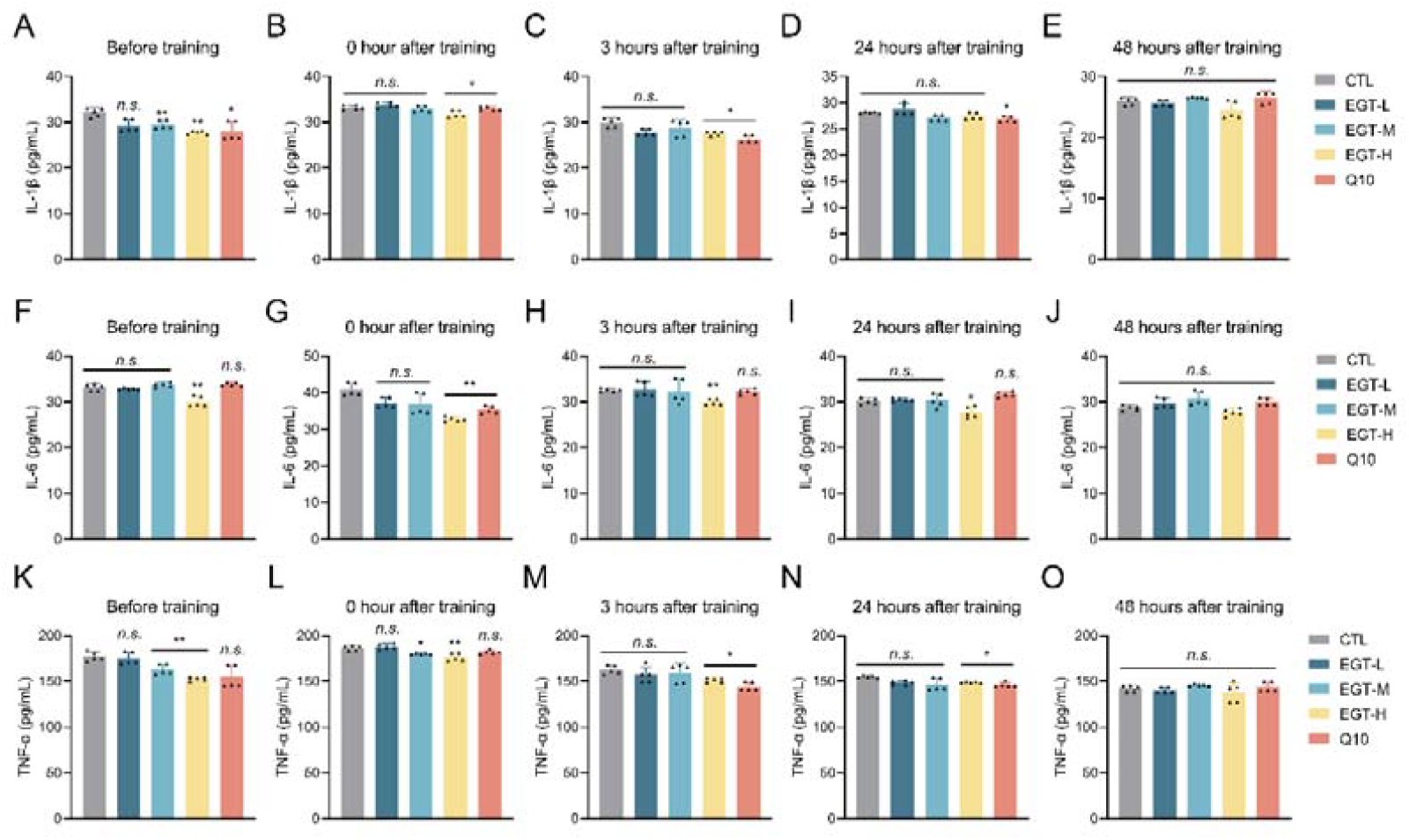
Effects of Gene III^**®**^ EGT on serum inflammatory cytokine levels in mice following the exhaustive exercise test. (A–E) Serum IL-1β (pg/mL): pre-training and 0 h, 3 h, 24 h, and 48 h post-training; (F–J) Serum IL-6 (pg/mL): pre-training and 0 h, 3 h, 24 h, and 48 h post-training; (K–O) Serum TNF-α (pg/mL): pre-training and 0 h, 3 h, 24 h, and 48 h post-training. The initial sample size was 10 animals per group; during detection, serum samples within the same group were pooled, resulting in 5 pooled samples per group for analysis (data points in the figure represent pooled samples). Data are expressed as mean ± SD. CTL: control group; EGT-L, EGT-M, and EGT-H: Gene III^**®**^ EGT groups at 10, 30, and 50 mg/kg, respectively; Q10: Coenzyme Q10 (50 mg/kg) positive control group.

As shown in the serum IL-6 results (Figure 3F–J), before training (F), only the EGT-H group was significantly lower than the CTL group, with a reduction of 9.39%. Immediately after training (0 h; G), the EGT-H and Q10 groups were significantly lower than the CTL group by 19.71% and 13.64%, respectively. At 3 h (H) and 24 h (I) post-training, only the EGT-H group was significantly lower than the CTL group, with reductions of 8.01% and 8.01%, respectively. No significant differences were observed at 48 h (J). This suggests that the IL-6-lowering effect of high-dose Gene III^**®**^ EGT is primarily manifested in the pre-training phase and the early post-exercise window up to 24 h.

As shown in the serum TNF-α results (Figure 3K–O), before training (K), the EGT-M and EGT-H groups were significantly lower than the CTL group by 7.84% and 13.65%, respectively. Immediately after training (0 h; L), the EGT-M group was 3.37% lower than the CTL group, while the EGT-H group showed a more significant reduction of 5.33%. At 3 h (M) and 24 h (N) post-training, the EGT-H and Q10 groups were significantly lower than the CTL group. At 3 h, they decreased by 6.74% and 11.20% compared to the CTL group, respectively; at 24 h, they decreased by 4.02% and 5.82%, respectively. No significant differences were observed at 48 h (O).

In summary, Gene III^**®**^ EGT intervention can reduce the levels of pro-inflammatory cytokines related to exhaustive exercise within different time windows. This suggests an overall palliative effect on exercise-induced inflammatory responses, potentially contributing to alleviating the post-exercise inflammatory burden and promoting recovery. Notably, the high-dose group demonstrated more prominent effects on IL-6 and, at certain time points, on IL-1β and TNF-α.

### 3.4 Gene III^®^ EGT Enhances Antioxidant Capacity and Reduces Oxidative Stress

The results of serum SOD activity (Figure 4A–E) showed that before training (A), the difference between the EGT-L group and the CTL group was not significant, while the EGT-M, EGT-H, and Q10 groups were significantly lower than the CTL group by 19.59%, 21.04%, and 22.76%, respectively. Immediately after training (0 h; B), SOD activity in all GENEIII ergothioneine dose groups (EGT-L, EGT-M, EGT-H) and the Q10 group was significantly lower than in the CTL group, decreasing by 12.72%, 13.90%, 11.55%, and 8.52% compared to the CTL group, respectively. At 3 h post-training (C), only the EGT-H and Q10 groups were significantly lower than the CTL group, with reductions of 20.28% and 19.50%, respectively. At 24 h post-training (D), all Gene III^®^ EGT dose groups were significantly lower than the CTL group; notably, the EGT-M group showed a more significant reduction. The groups decreased by 4.56%, 12.32%, and 14.98%, respectively. At 48 h post-training (E), all Gene III^®^ EGT dose groups and the Q10 group were significantly lower than the CTL group, decreasing by 10.35%, 5.85%, 7.84%, and 6.91% compared to the CTL group, respectively.

**Figure 4.**
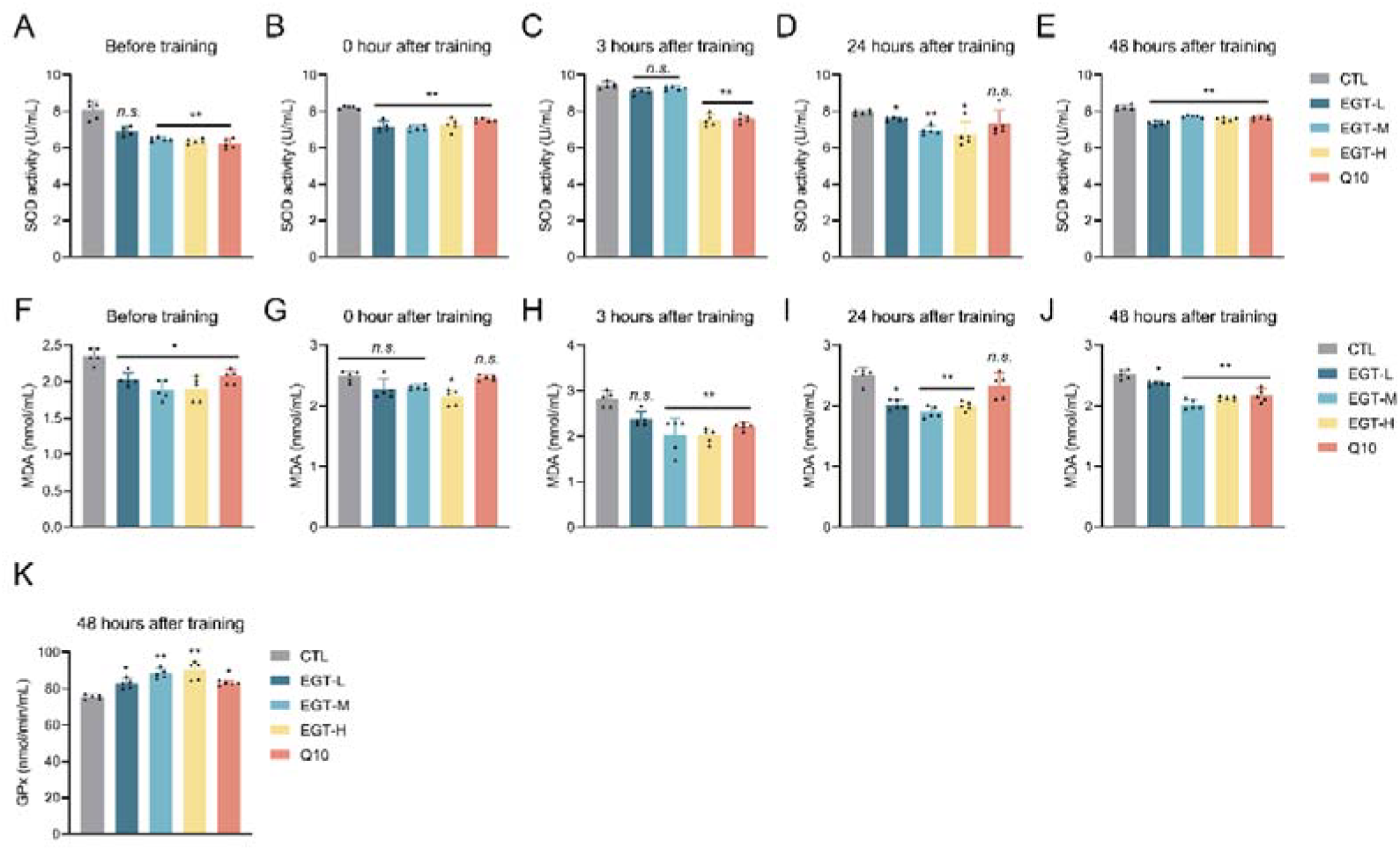
Effects of Gene III^®^ EGT on antioxidant-related markers in mice following the exhaustive exercise test. (A–E) Serum SOD activity (U/mL): pre-training and 0 h, 3 h, 24 h, and 48 h post-training; (F–J) Serum MDA (nmol/mL): pre-training and 0 h, 3 h, 24 h, and 48 h post-training; (K) Serum GPx activity (nmol/min/mL): 48 h post-training. The initial sample size was 10 animals per group; for analysis, serum samples within the same group were pooled in pairs, resulting in 5 pooled samples per group for detection and statistical analysis (data points in the figure represent pooled samples). CTL: control group; EGT-L, EGT-M, and EGT-H: Gene III® EGT groups at 10, 30, and 50 mg/kg, respectively; Q10: Coenzyme Q10 (50 mg/kg) positive control group.

As shown in the results for the lipid peroxidation marker MDA (Figure 4F–J), before training (F), all GENEIII ergothioneine dose groups and the Q10 group were significantly lower than the CTL group, decreasing by 13.74%, 19.83%, 18.89%, and 12.18%, respectively. Immediately after training (0 h; G), all Gene III^®^ EGT dose groups and the Q10 group were significantly lower than the CTL group, decreasing by 8.57%, 7.1%, 13.81%, and 0.94% compared to the CTL group, respectively. At 3 h post-training (H), the EGT-M, EGT-H, and Q10 groups were significantly lower than the CTL group, decreasing by 28.45%, 28.69%, and 21.37%, respectively, while the difference in the EGT-L group was not significant. At 24 h post-training (I), the EGT-L group was significantly lower than the CTL group, and the EGT-M and EGT-H groups showed even more significant reductions; the groups decreased by 19.17%, 24.1%, and 19.97%, respectively, while the difference in the Q10 group was not significant. At 48 h post-training (J), the EGT-L group was significantly lower than the CTL group, and the EGT-M, EGT-H, and Q10 groups showed even more significant reductions compared to the CTL group, decreasing by 5.95%, 19.58%, 15.74%, and 13.62%, respectively.

Additionally, the results of GPx activity detected during the recovery period at 48 h (Figure 4K) showed that the EGT-L and Q10 groups were significantly higher than the CTL group, and the EGT-M and EGT-H groups showed even more significant increases. Compared to the CTL group, they increased by 9.88%, 17.44%, 19.46%, and 10.01%, respectively.

In summary, Gene III^®^ EGT intervention can reduce MDA levels across multiple time windows after exhaustive exercise and increase GPx activity during the recovery period. This suggests that it helps alleviate exercise-induced lipid peroxidation damage and enhance the body’s antioxidant defense capacity. The changes in SOD activity at multiple time points also suggest corresponding regulation of the antioxidant system, which is consistent overall with the direction of “reducing oxidative damage and promoting recovery.”

### 3.5 Gene III^®^ EGT Modulates the AMPK/PGC-1α Signaling Pathway and Apoptosis in Skeletal Muscle

Following exhaustive exercise, the depletion of skeletal muscle energy substrate reserves, remodeling of mitochondrial energy metabolism pathways, and fluctuations in cell stress/apoptosis signaling are common molecular events in models of exercise fatigue and injury. Therefore, this study conducted a comprehensive evaluation during the recovery period (48 h) from three aspects: histology, energy metabolism signaling, and apoptosis-related molecules.

First, as skeletal muscle glycogen is an important energy substrate for endurance exercise, its post-exercise recovery can be used to reflect the overall state of energy substrate replenishment. Therefore, this study employed glycogen staining to observe the skeletal muscle of each group during the 48h recovery period. The results showed (Figure 5A) that the overall morphological structure of muscle fibers in each group was intact, and the glycogen staining intensity and distribution patterns were generally similar, with no obvious differences between groups. This suggests that under the detection time point and observation scale of this study, the effect of Gene III^®^ EGT intervention on skeletal muscle glycogen recovery was not significant or the difference was minor.

**Figure 5.**
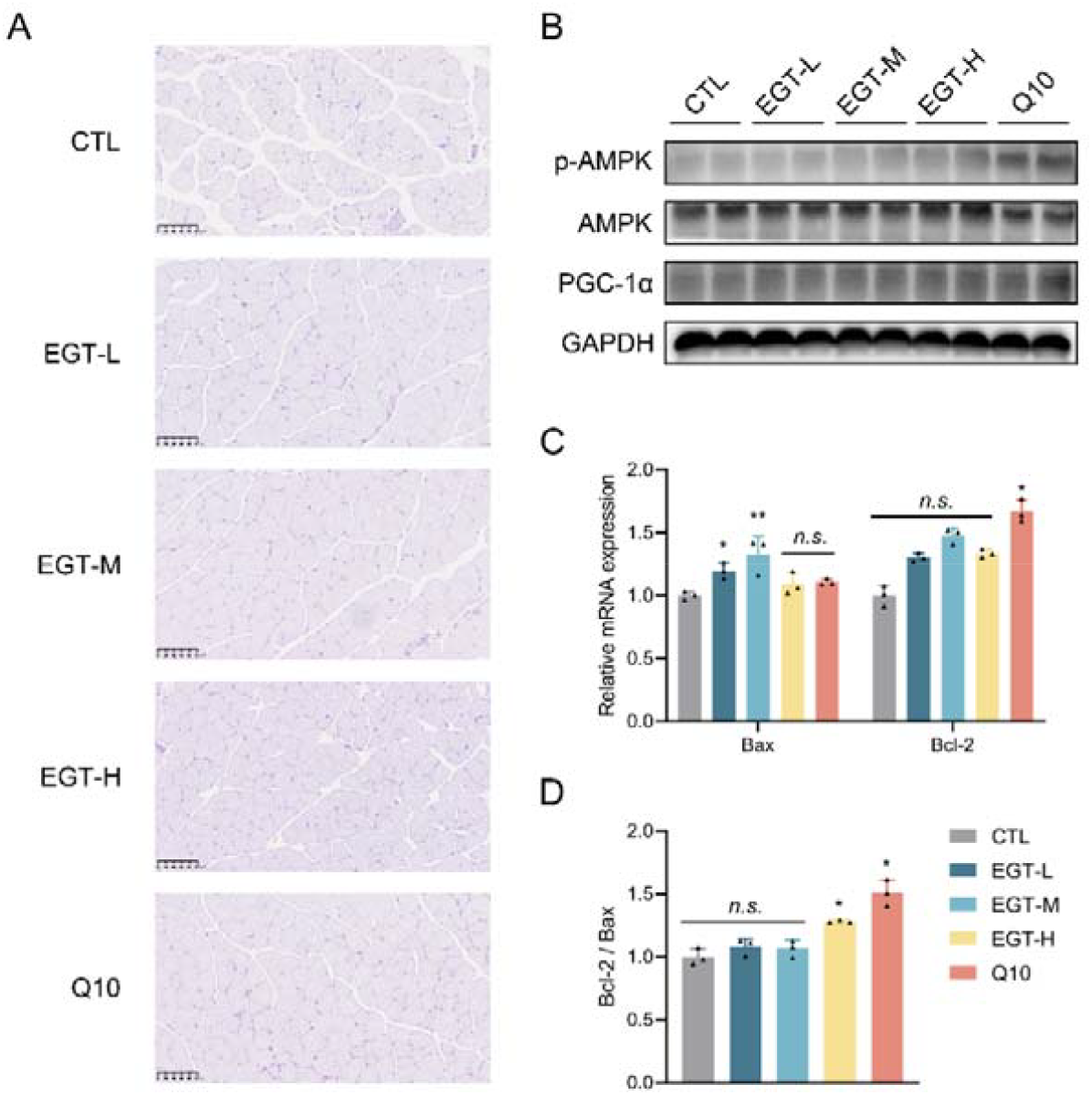
Effects of Gene III® EGT on skeletal muscle molecular markers following the exhaustive exercise test (48 h). (A) Representative images of glycogen-stained sections of pooled skeletal muscle tissue at 48 h (shown for groups: CTL, EGT-L, EGT-M, EGT-H, and Q10); (B) Representative Western blots showing protein expression of p-AMPK, AMPK, and PGC-1α in skeletal muscle, with GAPDH used as a loading control; (C) Relative mRNA expression levels of Bax and Bcl-2 (RT-qPCR), with 36B4 used as an internal control; (D) Bcl-2/Bax ratio. The initial sample size was 10 animals per group. For Western blot and RT-qPCR analyses, samples from the 10 animals within each group were pooled in equal amounts to generate one pooled sample per group. Three technical replicates (n=3) were performed for plotting and statistical analysis. Data are expressed as mean ± SD. CTL: control group; EGT-L, EGT-M, and EGT-H: GENEIII ergothioneine groups at 10, 30, and 50 mg/kg, respectively; Q10: Coenzyme Q10 (50 mg/kg) positive control group.

Secondly, AMPK is a skeletal muscle energy sensor; exercise and energy stress can promote AMPK phosphorylation activation (p-AMPK), thereby regulating energy metabolism and mitochondrial biogenesis. PGC-1α is a key transcriptional coactivator for mitochondrial biogenesis and enhanced oxidative metabolism, and its upregulation typically indicates improved metabolic adaptation and recovery capacity. Western blot results (Figure 5B) showed that compared with the CTL group, the p-AMPK and PGC-1α bands in the Gene III^®^ EGT intervention groups and the Q10 group showed an overall increasing trend, while changes in total AMPK were relatively less obvious. This suggests that the interventions may promote AMPK pathway activation and mitochondrial metabolism-related regulation.

Finally, Bax (pro-apoptotic) and Bcl-2 (anti-apoptotic) are a classic molecular combination reflecting post-exercise muscle cell stress and apoptotic tendency. An elevated Bcl-2/Bax ratio usually indicates that cells are in a more “anti-apoptotic/protective” state. RT-qPCR results (Figure 5C) showed that compared with the CTL group, Bax mRNA expression in the EGT-L group significantly increased, and the EGT-M group showed a more significant increase, rising by 20.61% and 38.52%, respectively, while differences in the EGT-H and Q10 groups were not significant. Regarding Bcl-2 mRNA, differences between the EGT-L, EGT-M, EGT-H groups and the CTL group were not significant, while the Q10 group significantly increased by 66.82%. Further calculation of the Bcl-2/Bax ratio (Figure 5D) revealed that differences between the EGT-L and EGT-M groups and the CTL group were not significant, while the Bcl-2/Bax ratio in the EGT-H and Q10 groups was significantly higher than in the CTL group, increasing by 28.17% and 50.9%, respectively. This suggests that high-dose GENEIII ergothioneine and the positive control tend to improve the balance of apoptosis-related molecules and enhance the cellular protective state during the recovery period.

In summary, Figure 5 suggests from the aspects of glycogen reserves, AMPK–PGC-1α energy metabolism signaling, and Bcl-2/Bax apoptotic balance that Gene III^®^ EGT may facilitate tissue recovery and functional reconstruction after exhaustive exercise by promoting metabolic pathway adaptation and improving the state of cell stress-related molecules during the recovery period.

## 4. Discussion

The primary objective of this study was to evaluate the anti-fatigue efficacy of Gene III^®^ Ergothioneine (EGT) and to elucidate its molecular mechanisms in a mouse model of exhaustive exercise. Our findings demonstrate that EGT supplementation, particularly at medium and high doses, significantly extends time to exhaustion and running distance, with efficacy comparable to the established supplement Coenzyme Q10. This improvement in endurance performance was substantiated by a marked reduction in serum metabolic byproducts. Specifically, Gene III^®^ EGT lowered lactic acid (LA) and blood urea nitrogen (BUN) levels immediately post-exercise and during recovery. The accumulation of lactate is a classic marker of anaerobic glycolysis that inhibits muscle contraction, while elevated BUN indicates increased protein catabolism. The reduction in these markers, alongside decreased LDH levels, suggests that EGT enhances aerobic metabolic efficiency and preserves muscle cell membrane integrity under stress.

Mechanistically, the anti-fatigue effects of Gene III^®^ EGT appear to be mediated through a dual regulation of oxidative stress and inflammation. Intense exercise overwhelms endogenous antioxidant defenses, leading to lipid peroxidation and the release of pro-inflammatory cytokines, which exacerbate central and peripheral fatigue. In this study, EGT significantly reduced MDA levels and pro-inflammatory cytokines (IL-1β, IL-6, TNF-α) while upregulating SOD and GSH-Px activities. Unlike common antioxidants, EGT is actively transported and accumulated in tissues with high oxidative stress via the specific transporter OCTN1. This specific retention likely explains the sustained antioxidant effects observed up to 48 hours post-exercise. By neutralizing ROS and blunting the inflammatory surge, EGT likely interrupts the feedback loop that signals fatigue to the central nervous system.

A novel highlight of this study is the demonstration that Gene III^®^ EGT modulates the AMPK/PGC-1α signaling pathway in skeletal muscle. AMPK acts as a cellular energy sensor; its phosphorylation triggers PGC-1α, the master regulator of mitochondrial biogenesis and oxidative metabolism. We observed that EGT treatment enhanced the expression of p-AMPK and PGC-1α, suggesting that EGT functions not merely as a radical scavenger but as a signaling modulator that improves mitochondrial adaptability. Furthermore, EGT increased the Bcl-2/Bax ratio, shifting the cellular balance towards survival and protecting muscle cells from exercise-induced apoptosis[19].

Despite these promising findings, this study has limitations. We focused on the gastrocnemius muscle without distinguishing between oxidative (Type I) and glycolytic (Type II) fiber types, which may exhibit different OCTN1 expression profiles. Additionally, while we identified the involvement of the AMPK pathway, specific inhibitors were not used to confirm causality. Future research should address these aspects and translate these murine findings to human physiology. In conclusion, Gene III^®^ EGT effectively alleviates exercise-induced fatigue by mitigating metabolic stress and inflammation, and notably, by activating the AMPK/PGC-1α pathway to promote mitochondrial function and cellular protection. These results highlight the potential of Gene III^®^ EGT as a nutritional supplement for fatigue management.

